# Sticking around: Optimal cell adhesion patterning for energy minimization and substrate mechanosensing

**DOI:** 10.1101/2020.08.17.253609

**Authors:** Josephine Solowiej-Wedderburn, Carina M. Dunlop

**Affiliations:** Department of Mathematics, University of Surrey, Guildford, UK; Centre for Mathematical and Computational Biology and Department of Mathematics, University of Surrey, Guildford, UK

## Abstract

Cell mechanotransduction, in which cells sense and respond to the physical properties of their micro-environments, is proving fundamental to understanding cellular behaviours across biology. Tissue stiffness (Young’s modulus) is typically regarded as the key control parameter and bioengineered gels with defined mechanical properties have become an essential part of the toolkit for interrogating mechanotransduction. We here, however, show using a mechanical cell model that the effective substrate stiffness experienced by a cell depends not just on the engineered mechanical properties of the substrate but critically also on the particular arrangement of adhesions between cell and substrate. In particular, we find that cells with different adhesion patterns can experience two different gel stiffnesses as equivalent and will generate the same mean cell deformations. For small adhesive patches, which mimic experimentally observed focal adhesions, we demonstrate that the observed dynamics of adhesion growth and elongation can be explained by energy considerations. Significantly we show different focal adhesions dynamics for soft and stiff substrates with focal adhesion growth not preferred on soft substrates consistent with reported dynamics. Equally, fewer and larger adhesions are predicted to be preferred over more and smaller, an effect enhanced by random spot placing with the simulations predicting qualitatively realistic cell shapes in this case. The model is based on a continuum elasticity description of the cell and substrate system, with an active stress component capturing cellular contractility. This work demonstrates the necessity of considering the whole cell-substrate system, including the patterning of adhesion, when investigating cell stiffness sensing, with implications for mechanotransductive control in biophysics and tissue engineering.

**Author summary:** Cells are now known to sense the mechanical properties of their tissue micro-environments and use this as a signal to control a range of behaviours. Experimentally, such cell mechanotransduction is mostly investigated using carefully engineered gel substrates with defined stiffness. Here we show, using a model integrating active cellular contractility with continuum mechanics, that the way in which a cell senses its environment depends critically not just on the stiffness of the gel but also on the spatial patterning of adhesion sites. In this way, two gels of substantially different stiffnesses can be experienced by the cell as similar, if the adhesions are located differently. Exploiting this insight, we demonstrate that it is energetically favourable for small adhesions to grow and elongate on stiff substrates but that this is not the case on soft substrates. This is consistent with experimental observations that nascent adhesions only mature to stable focal adhesion (FA) sites on stiff substrates where they also grow and elongate. These focal adhesions (FAs) have been the focus of work on mechanotransduction. However, our paper demonstrates that there is a fundamental need to consider the combined cell and micro-environment system moving beyond a focus on individual FAs.

## Introduction

It is becoming increasingly apparent that mechanical cues have an impact on cellular behaviour affecting, for example, the growth, differentiation and ultimate fate of cells [1–5]. Considering stem cells in particular, the rigidity of the substrate on which they are cultivated has been shown to determine their specialisation, overriding any additional chemical cues over an extended period of weeks or even without their input entirely [6,7]. Throughout the body, cells are influenced by the mechanical properties of their surroundings, for example, recent studies have linked osteoporosis with mechanical changes in bone tissue [8] and the ageing of stem cells within the central nervous system with the stiffening of their microenvironment [9].

Experimental investigations of mechanotransduction commonly focus on stiffness as a single control parameter. This has stimulated activity in developing biomaterials with defined stiffness, ligand density and functionalisation [10–12]. As well as the changes in behaviours mentioned, several structural changes have been identified that occur in response to changes to substrate stiffness. Cell shape is observed to be altered on soft versus stiff substrates, with cells adopting smaller, rounder shapes on softer substrates and appearing larger and more angular on stiffer substrates [13–17]. Equally the distribution of adhesion sites and their size has been found to be dependent on gel stiffness. For example, finding that the size of focal adhesions (FAs) increase on stiffer substrates [14,16,18,19], whereas on soft substrates nascent adhesions do not stabilise into FAs [15,20]. The effect of cell shape and geometric constraints on cell behaviours have been investigated using micro-patterning techniques [21]. In these studies, surface functionalisation is used to constrain cell adhesion to pre-defined regions [22]. A significant set of studies have focused on areas of complete adhesion in specific geometries (e.g. circular, triangular, square) [23, 24], these have demonstrated that adhered area and shape can control a range of cellular behaviours including proliferation and signalling [5, 22, 25]. Further studies have used micropatterning to uncouple the effects of adhesion size and cell spreading [26, 27].

As microscopy and biophysical tools advance, we are beginning to understand the structure of cells and their ability to generate forces [28]. Central to this and much studied is the integrin binding of cells to the extracellular matrix (ECM). A particular focus of mechanobiology has been the FA complexes, which generate small patches of strong attachment between the cell and the gel on stiffer substrates. (For details of the structure of the cytoskeleton and FA dynamics see the reviews [20,29–31]). Cells generate active contractile forces while attached to the underlying substrate. These forces are generated by myosin motors and are transmitted to the ECM through the adhesions. The substrate resistance experienced consequently allows the cell to mechanically sense its environment although the details of the physical mechanism of this signal transduction remains unclear. FAs have received most attention as mechanosensors [32–34], although there is an increasing awareness of the need to account for mechanosensing across the cell including at the nuclear envelope [35, 36].

Several theoretical models have been developed to gain an insight into the cellular force generation and its effects. Largely, these treat the main body of a cell as an elastic solid which is being acted upon by an active component; this is coupled to a substrate which offers further resistance to the force. The way in which the active cellular contractility is represented broadly falls into two categories: simulations of cytoskeletal dynamics and active continuum theories. Computational simulations of the cytoskeleton tend to focus on the dynamics of subcellular constituents of the contractile mechanism to investigate the cell-scale effects of their collective behaviour (e.g. [37–39]). In the continuum approach, an adaptation of linear elasticity can lead to the introduction of an active contractile term to the material constitutive relations and in this way either a force balance equation [40,41] or an equivalent energy minimisation [42] problem can be constructed for the cell deformations and stress. Such models can be adapted to incorporate different adhesion dynamics, including FA clustering [43], to investigate intracellular mechanics [16].

In this work, we adopt an active stress approach to model the cellular contractility. We focus on the significance of adhesion distribution and patterning on a cell’s overall ability to deform. Two distinct cases are considered. First, we consider that the cell is adhered in a ring around its edge before considering the case that the cell is adhered at several distinct spots mimicking FA complexes. In the case of spot adhesion we vary the distribution, total area and shape of adhered regions and consider the effects on the mean cellular deformation, relating this to the resistance the cell experiences from the underlying substrate. These results show that the substrate may be experienced as more or less stiff depending on how the cell is adhered. Specifically, we show that a cell with a sparse distribution of adhered regions around its periphery, with large gaps between them, effectively experiences a softer substrate than a cell with more continuously distributed adhesions around its edge. Cell morphology is also observed to change with stiffness qualitatively agreeing with experimentally observed shape changes when the adhesion points are randomly distributed. Energy calculations additionally show that it is energetically favourable, reducing the work done to the substrate, for the cell to be adhered at spots with large variance in inter-spot spacing. Significantly, on stiff substrates growth and elongation of these adhesion patches is energetically favoured whereas on soft substrates this is not the case.

## Results

### Model

We use the active stress formulation for modelling contractile cells on soft gel substrates [26,40,42,44–46]. Thus the cell stress *σ* = *σ^P^* + *σ^A^*, where *σ^A^* is an active component of stress generated by the contractile machinery embedded in the cytoskeleton. We assume throughout that there is a constant target contraction *P*_0_, which is generated by action of the molecular motors within the acto-myosin network. We additionally assume linear elasticity of both cell and gel, noting that the timescale for cell adhesion is faster than the viscoelastic relaxation timescale [42], see also Methods. Cell adhesion to the underlying substrate is captured through the force balance (assuming plane stress)

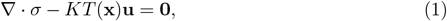

where the cell is not adhered *T*(**x**) = 0 and *T*(**x**) = 1 where the cell is adhered. In the case of uniform adhesion (*T*(**x**) ≡ 1) we recover the force balance considered in [40,46]. The parameter *K* describes the stiffness of the substrate, with *K* increasing with substrate stiffness. The parameter K may be related to the material properties of some common experimental systems, for example, for an array of micropillars of stiffness *k* and number density *N, K* = *Nk* [10,47] while for a thin elastic gel substrate with shear modulus *μ_S_* and thickness *h_S_* the rigidity parameter *K* ≈ *μ_s_/h_S_* [45,46]. We here consider two cases for *T*(*x*). The first is that the cell is completely adhered around its edge and the second that the cell is adhered at distinct spots that are spatially distributed around the edge. This latter case can be considered to describe focal adhesions (FAs). In the case of a ring

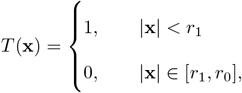

depicted in Fig 1A, and by symmetry the deformations are purely radial so that **u** = *u*(*r*)**e_r_**. Consequently, the force balance equation (1) can be solved analytically to give

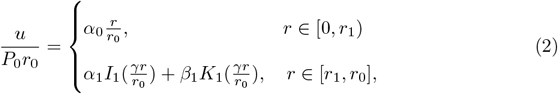

where *α*_0_, *α*_1_ and *β*_1_ are constants determined from the boundary conditions (see Methods) and *I*_1_ and *K*_1_ are modified Bessel functions. The contraction parameter *P*_0_ linearly scales the deformation altering its magnitude only. As an illustration, Fig 1B plots the analytical solution for different thicknesses of adhesive ring. We discover one key nondimensional parameter, 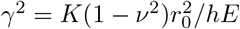 which quantifies the substrate resistance for a cell with Young’s modulus *E*, Poisson ratio *ν*, diameter 2*r*_0_ and thickness *h*. For a stiffer substrate (*K* larger) *γ* is greater.

**Fig 1.**
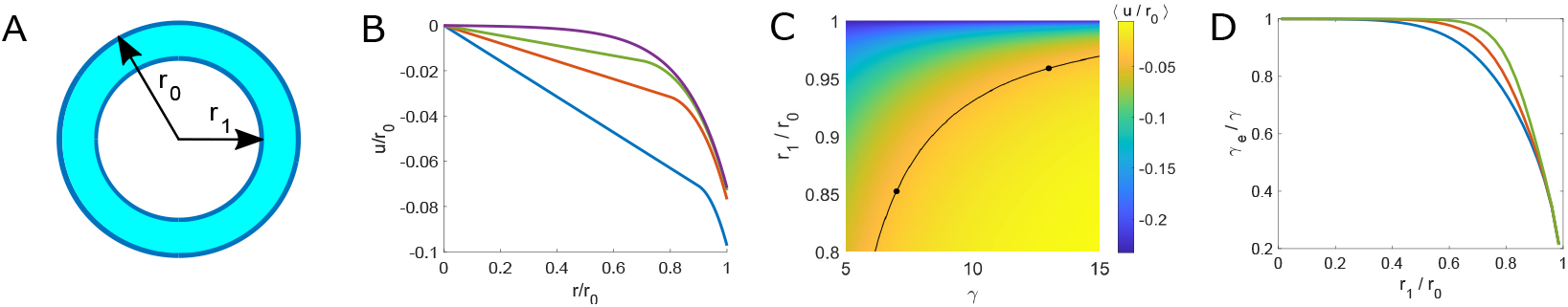
Adhesive ring thickness changes the effective substrate stiffness experienced. (A) Schematic diagram of circular cell with an adhesive ring. (B) Plot of deformation profile of a cell with *r*_1_/*r*_0_ = 0.9, 0.8, 0.7, 0 [from bottom to top]. *r*_1_/*r*_0_ = 0 corresponds to complete adhesion (*γ* = 7). (C) Heat map showing how mean cellular deformation varies with ring thickness (parameterized by *r*_1_) and substrate stiffness (parameterized by *γ*). Along the contour line (black) cells display the same mean deformation. (D) Relative effective resistance plotted against internal ring radius (*r*_1_) on substrates with *γ* = 5, 10, 15 [from bottom to top]. (Here and in all further figures we set *P*_0_ = 0.7 and *ν* = 0.45.)

To model localised spots of adhesion and in particular FAs, we take *T*(*x*) = 1 only in small circular or elliptical regions in the cell (e.g. Fig 2A). In this case, an analytical solution cannot be obtained and numerical solutions are obtained using finite element methods (see Methods), see as an example Fig 2. In the case of spots, *γ* remains the key control parameter quantifying the substrate resistance.

**Fig 2.**
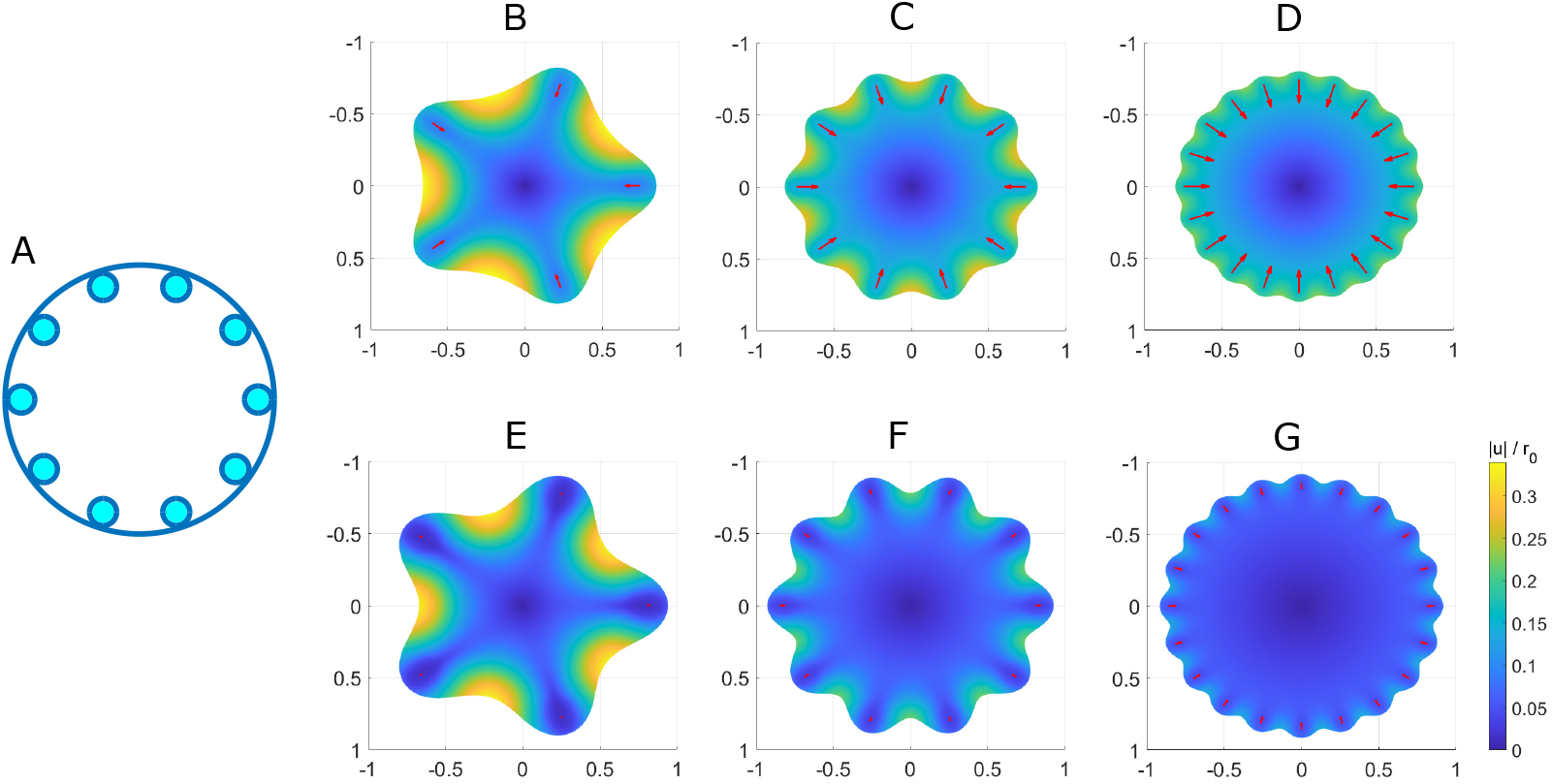
Arrangement of adhesion into localised spots facilitates localised regions of high deformation. (A) Schematic diagram of a circular cell with 10 spots evenly distributed around the edge. (B)-(E) heat maps of the mean deformation on soft substrates with *γ* = 5 (B)-(D) or stiff with *γ* = 11.34 (E)-(G) with arrangements of 5 (B and E), 10 (C and F) and 20 (D and G) spots evenly distributed around the cell edge. Red arrows show the deformation of the mid-point of each spot. Adhered area is 10%A, where *A* is the pre-contraction cell spread area (i.e. 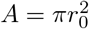).

### Adhesive ring thickness changes the effective substrate stiffness experienced

For cells adhered uniformly at their outer edge we observe in Fig 1B that the cell’s ability to deform is affected by the width of its adhesive ring, with thicker rings exhibiting reduced deformations. To better quantify the potential effect of adhered area on mechanosensing it is necessary to define a measurable quantity for comparison. One such measure adopted which is relatively easy to interpret is the mean deformation over the cell area [48]. Where the mean deformation is lower the apparent substrate stiffness by this measure would be higher, whereas a larger mean deformation would correspond to a softer substrate. For the adhered ring the mean cellular deformation (scaled by cell radius) can be explicitly calculated, see Methods.

Fig 1C shows how the mean cellular deformation varies with the adhesive ring width and substrate resistance parameter *γ*. Cells with the thinnest adhered rings and on the least resistant substrates display the greatest mean cellular deformation. Also plotted in Fig 1C is an illustrative contour along which the mean deformation is constant (〈*u*/*r*_0_〉 = −0.04). Specifically, it can thus be seen that a cell with *r*_1_/*r*_0_ ≈ 0.85 on a substrate *γ* = 7 experiences the same mean deformation as a cell with *r*_1_ /*r*_0_ ≈ 0.96 on a substrate *γ* = 13 and thus potentially senses both as equally stiff. Thus a cell with ring of adhesion thickness *r*_1_ /*r*_0_ = 0.85 on a substrate engineered at stiffness *γ* = 7 experiences this substrate as if it were of stiffness *γ* ≈ 4.6 (choosing as reference the state of full adhesion).

This observation informs our definition of an *effective substrate stiffness γ_e_* for the adhesive pattern, which is the *γ* on the contour of constant mean deformation corresponding to complete adhesion. *γ_e_* may be calculated by solving 〈*u*(*γ*)〉 = 〈*u_CD_*(*γ_e_*)〉 where *u_CD_* is the solution for a completely adhered disc. This solution is obtained numerically with a Newton-Raphson iterative scheme and plotted in Fig 1D. We see that throughout *γ_e_* < *γ*, so that the resistance experienced by the partially adhered cell is less than that for a completely adhered cell. Specifically, cells with thinner rings (with *r*_1_/*r*_0_ near 1) sense an effectively softer substrate as they experience less resistance, however, this effect becomes less pronounced for wider adhesive rings. For example when *r*_1_ ≈ 0.66*r*_0_, *γ_e_*/*γ* is already close to 1 (*γ*_e_/*γ* = 0.9 on *γ* = 5), showing that the cell is sensing almost the same resistance as it would when completely adhered. This effect is enhanced on stiffer substrates, for instance when *γ* = 15, *γ_e_*/*γ* = 0.9 is obtained at *r*_1_ ≈ 0.78*r*_0_, see S1 Fig.

Other measures that may correlate with mechanosensing demonstrate very similar heat maps to Fig 1C with reduced adhesion resulting in a lower effective stiffness with a similar dependency on substrate stiffness. This is to be expected given the close relationship between deformation, strain and energy arguments. See, for example, S2 Fig, where the measures based on, maximal cellular deformation, mean cellular strain and maximal cellular strain [44] are considered with energy arguments considered in a later section.

### Arrangement of adhesion into localised spots (FAs) further reduces the effective stiffness of the substrate

To consider the effect of breaking up cellular adhesion into small spots of adhesion (mimicking the focal adhesions) we introduce a distribution of circular spots placed around the cell periphery. The number of spots is then varied maintaining a constant adhered area, this increases the between spot gaps. The outputs of simulations with 20, 10, and 5 spots on soft and stiff substrates are shown in Figs 2B-G. The corresponding mean cellular deformations and effective resistance parameters are given in Table 1. It is clear that reducing the number of adhesions increases the gap size between adhered regions, thus facilitating localised areas of high deformation On the stiffer substrate the cellular deformations are as expected smaller. Where the gaps are larger, however, the effect of substrate stiffness is reduced. Considering, for example, the maximal deformation with 20 spots we find a 40% decrease in the maximum deformation from *γ* = 5 to *γ* = 11.34, while with 5 spots the decrease in maximum deformation is only 5% with the same increase in substrate stiffness.

**Table 1.** Arrangement of adhesion into localised spots reduces the effective stiffness of the substrate. Mean cellular deformation and corresponding effective resistance parameter for different spot distributions for two different *γ* corresponding to soft and stiff substrates. Other parameters as for Fig 2.

| | $\gamma = 5$ | | $\gamma = 11.34$ | |
| --- | --- | --- | --- | --- |
| | $\langle u/r_0 \rangle$ | $\gamma_e$ | $\langle u/r_0 \rangle$ | $\gamma_e$ |
| 20 spots | 0.133 | $0.37\gamma = 1.84$ | 0.055 | $0.33\gamma = 3.75$ |
| 10 spots | 0.144 | $0.33\gamma = 1.67$ | 0.084 | $0.25\gamma = 2.82$ |
| 5 spots | 0.166 | $0.27\gamma = 1.35$ | 0.133 | $0.16\gamma = 1.84$ |

In Table 1, we present the mean cellular deformations corresponding to the simulations in Fig 2. We see that on substrate *γ* = 5 halving the number of spots from 20 to 10 increases the mean deformation by 8.3%. Thus demonstrating that decreasing the number of adhered spots results in an increased mean cellular deformation so that by decreasing the number of adhered regions a cell effectively senses a softer substrate. This effect is enhanced on stiffer substrates, for example when *γ* = 11.34 halving the number of spots from 20 to 10 increases the mean deformation by 52.7%. Furthermore, a cell with 20 spots on substrate *γ* = 5 may effectively sense the same resistance (*γ_e_* = 1.84) as with 5 spots on substrate *γ* = 11.34. We reiterate that the adhered area is the same in each case and it is only the rearranging of the sites of cellular adhesion that enables a cell to experience this softer environment.

### Increasing spot size increases apparent substrate stiffness - an effect which may be compensated for by the elongation of spots into elliptical patches

Increasing individual spot size while keeping the number of adhesions fixed has the natural effect of increasing the adhered area. As such we find that the mean cellular deformation decreases with the increased resistance from a greater adhered area Fig 3A. Correspondingly, we suggest that the cell effectively experiences a stiffer substrate with an increase in adhered area, in analogy to the results for a cell with an adhesive ring. However, this effect can in part be compensated for if the increase in area is not uniform but is generated through an elongation of the spot. In Fig 3B, we show how elliptical spots affect the mean deformation. Increasing the aspect ratio of the spots (for a given spot size), we find that elongating the spots inwards may result in an increase in the mean cellular deformation (Fig 3B) potentially compensating for the reduction the increased area has imposed. For example, in Fig 3B for a cell with 10 spots we see that increasing the adhered area from 10% to 12% without altering the spot shape would decrease the mean deformation substantially, however, by increasing the the spot aspect ratio from *b/a* = 1.15 to *b/a* = 5.89 the mean deformation can be kept constant. The mean cellular deformation can similarly be conserved when increasing the adhered area to 14% with a spot aspect ratio of *b/a* = 8.44. Experimentally it is observed that FAs grow as the applied force increases, but that they additionally tend to elongate in the direction of applied forces [19,20]. Our results suggest that that such an elongation may be being used to at least partially compensate for the effect of adhesion growth.

**Fig 3.**
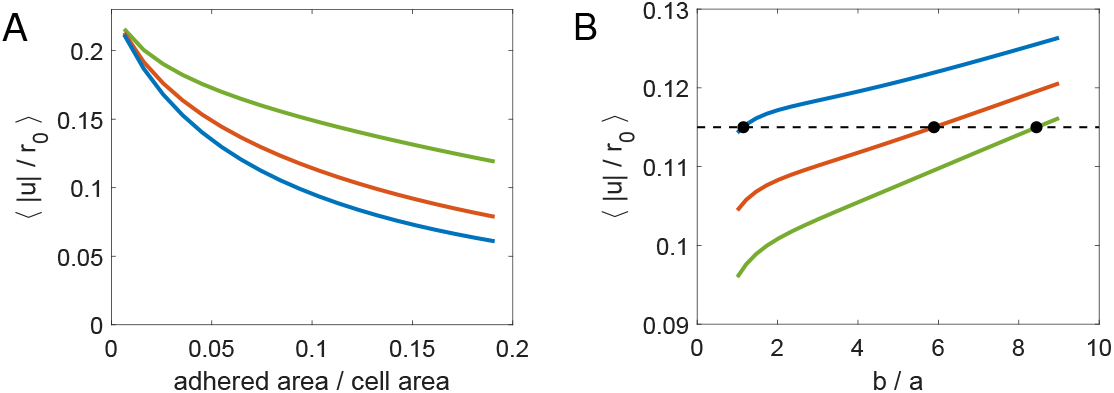
Increasing spot size increases apparent substrate stiffness: an effect which may be compensated for by the elongation of spots into elliptical patches. (A) Mean deformation plotted against adhered area for evenly distributed adhesive spots, with number of spots 5, 10, 20 [from top to bottom]. (B) Mean deformation plotted against spot aspect ratio for 10 evenly distributed spots with adhered area *A*_ad_ = 10%, 12%, 14% [from top to bottom]. Increasing the aspect ratio *b/a* corresponds to an elongation towards the centre of the cell. The black dotted line indicates 〈|*u*|/*r*_0_〉 = 0.115. (In (A) and (B) *γ* = 7.)

### Focal adhesion growth reduces strain energy on stiff substrates but not on soft substrates; elongated adhesion spots are energetically favourable

Considering now the strain energy *W*, of the system, this can be expressed as *W* = *W_CA_* + *W_CP_* + *W_S_*, where *W_CA_* is the work done by the active contractile network of the cell, *W_CP_* the strain energy in the passive cell components and *W_S_* the substrate strain energy (see Methods for the integral formulations of these energies). As this is a closed system, we expect no net loss or gain of energy and so *W* = 0. The work done *W_CA_* thus causes both the deformation of the cell and surrounding substrate and is equal to −(*W_CP_* + *W_S_*). For constant cellular contractility the active work done by the cell is directly proportional to its mean strain (see Methods). Thus, the behaviour of *W_CA_* is qualitatively very similar to the mean cellular deformation both for ring adhesion and adhesive spots as discussed above, see S3 Fig. We focus here on the substrate strain energy *W_S_*, which is often experimentally used to quantify the mechanical activity of a cell and its contractile strength [49–51]. Indeed *W_S_* has recently been identified as an important metric to describe the entire output work done across different cell types, morphologies and substrates [52].

In Figs 4A-E we consider how the adhered area affects *W_S_* in particular focusing on the case where spot radius is increased. (For the case of an increasingly wide ring of adhesion see S4 Fig). In Fig 4D and E we consider the particular cases of a cell adhered to a soft substrate (*γ* = 5) and a stiff substrate (*γ* = 15). For comparison we plot the analytical solution for a completely adhered ring of the same area. We observe that, as expected, the work done to the substrate by cells with adhered spots tends towards the continuous solution of an adhesive ring as the spot distribution becomes more dense. Similarly when there are fewer adhered regions the cell does less work to the substrate. We observe that on soft substrates as the adhered area increases (spot radius increases) so the substrate energy increases making this energetically unfavourable, although there is a turning point point in this behaviour. By contrast on the stiff substrate it is energetically favourable to increase adhesion size in all of the spot distributions considered. This can be explained as on stiff substrates increasing adhesion reduces the potential cell deformation and substrate deformation as the cell is now fixed in place due to the rigidity of the substrate. However, in the case of 5 spots we see a turning point in this behaviour, beyond which *W_S_* increases as adhesions continue to grow.

**Fig 4.**
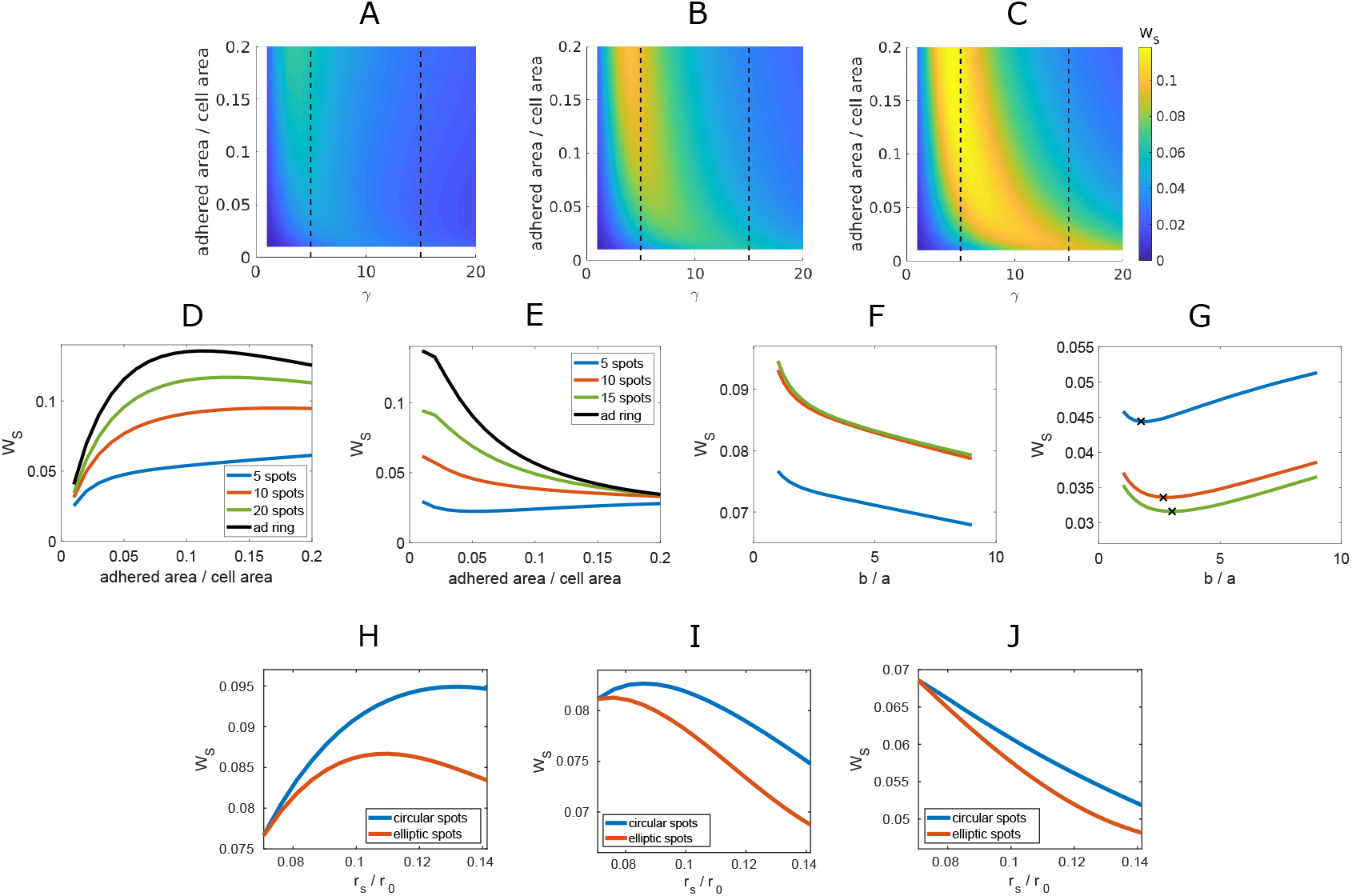
Focal adhesion growth reduces strain energy on stiff substrates but not on soft with spot elongation optimal. (A)-(C) heat maps showing substrate strain energy against *γ* and the proportion of adhered area with 5 (A), 10 (B) and 20 (C) adhered spots. (D),(E) *W_S_* against adhered area for for *γ* = 5 (D) and *γ* =15 (E), as indicated by the dotted lines in (A)-(C), for 5, 10 and 20 spots. *W_S_* for an adhered ring plotted for comparison. (F),(G) *W_S_* against spot aspect ratio (*b/a*) for *γ* = 5 (F) and *γ* =15 (G) for 10 spots at adhesion 5%, 10%, 15% in blue, orange and green, respectively. (H)-(J) *W_S_* for an even distribution of 10 spots on substrates with *γ* = 5 (H), *γ* = 7 (I) and *γ* =10 (J). Blue line indicates circular spots of spot radius *r_s_*. Orange line corresponds to elliptical spots with a fixed width but increasing length so the aspect ratio increases as adhered area increases, here *W_S_* is plotted against the equivalent radius of circular spots.

Finally, we consider the effects of an elongating spot on the substrate strain energy, shown in Figs 4F and G. In Fig 4F, we see that on a soft substrate increasing the aspect ratio from circular to elongated elliptic spots (aligned towards the cell centre) decreases the work done to the substrate as the cell is more able to deform in the gaps between adhesions. Furthermore, our investigations suggest that on stiffer substrates there is an optimal spot aspect ratio for these adhesions depending on spot size. In the case of an adhered area ranging from 5% to 15% and *γ* =15 this is approximately 2-3 times as long as they are wide (Fig 4G). In Figs 4H-J we consider the effects of increasing the adhered area. Comparing the effect of increasing the radius of circular spots, to maintaining a fixed width and accommodating the extra area by elongating the spots towards the centre of the cell we see that elliptic spots result in a lower *W_S_*. This combined suggests the elongation of adhesions is energetically favourable.

### Random placement of adhesion sites can generate apparently softer substrates compared with uniform placement and is energetically favourable

To investigate the effect of more realistic distributions of adhesion in which spots are distributed non-uniformly we considered two cases. In the first, spots are distributed at the cell edge but at a random angle, see e.g. Figs 5C and E. In the second, spots are distributed randomly within an annular region located near the edge of the cell, see e.g. Figs 5D and F. In all cases, adhesion is considered to localise at the cell edge as is experimentally observed. We first note that the predicted cell shapes (with both arrangements) are now qualitatively very similar to those observed experimentally.

**Fig 5.**
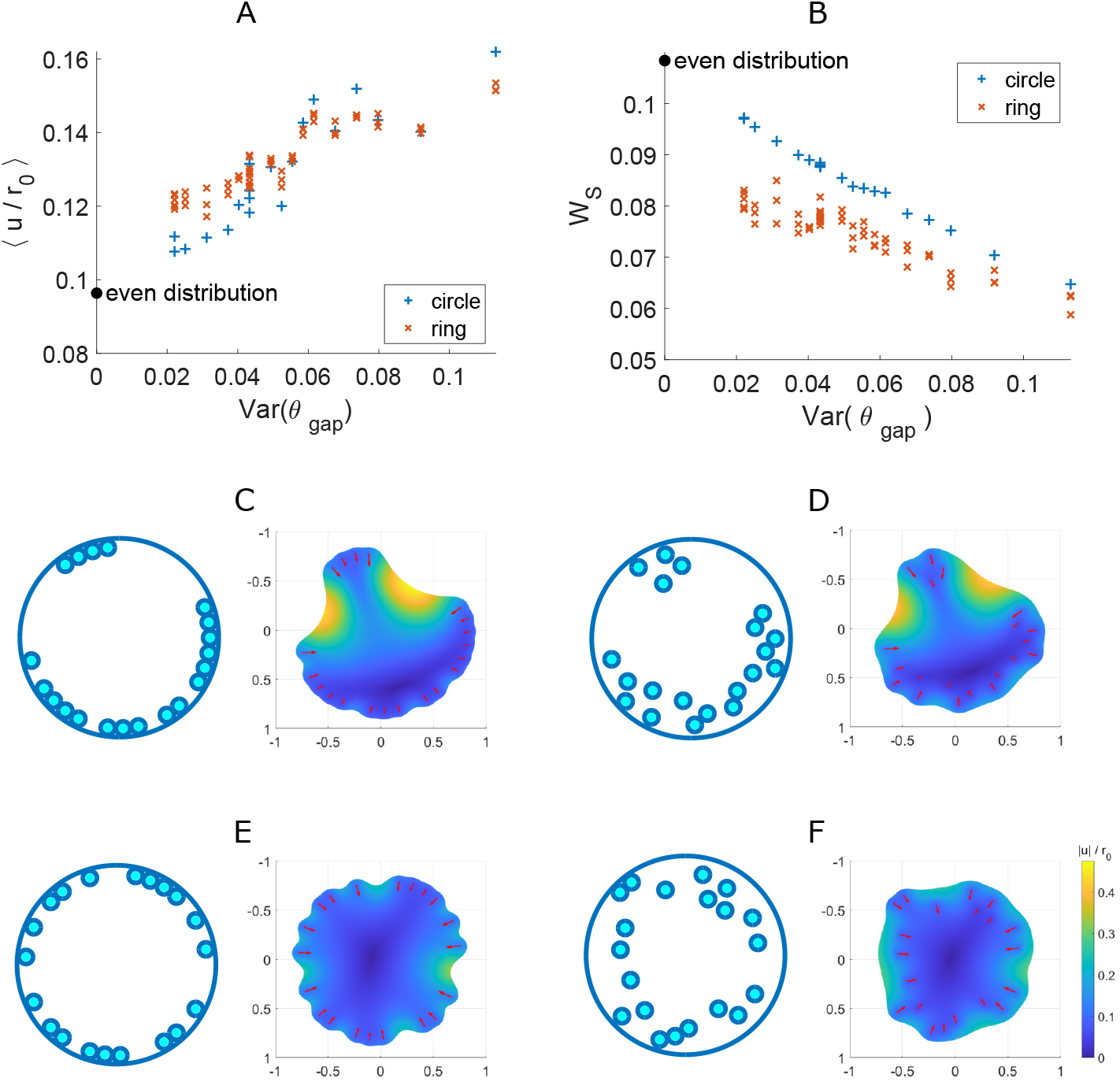
Random placement of adhesion sites can generate apparently softer substrates compared with uniform placement and is energetically favourable. (A) Mean cellular deformation and (B) substrate strain energy plotted against the variance in angular gap size for spots restricted to the edge (blue pluses) and in an annular region (0.6*r*_0_ < *r* < *r*_0_) (orange crosses). Results for an even distribution of spots are included for comparison at Var(*θ_g_*) = 0. (In each simulation there are 20 circular spots, covering 10% of the cell spread area, for the ring the same angular spot placements are chosen but the radial position is varied; *γ* = 7). (C)-(F) examples of the cell deformation observed in the above simulations. (C) corresponds to point in (A) with largest mean deformation; (D) is a corresponding ring distribution. (E) corresponds to the point in (A) with the least mean deformation; (F) is a corresponding ring distribution.

In Figs 5C and E, we show the deformation of cells with spot distributions with the greatest and least variance in angles (*θ_g_*) between adjacent spots from the multiple simulations run for each arrangement. Where Var(*θ_g_*) is larger we observe few larger clusters and the existence of larger gaps between clusters, this results in greater mean cellular deformation, see Fig 5A. In Fig 5B we see that the substrate strain energy is less where Var(*θ_g_*) is larger. This corresponds to the result we found in Figs 4D and E.

To compare the two arrangements (adhesions at the cell edge as compared to within a constrained ring) spot distributions within the ring were chosen to have the same angular distribution (with different radial positions) as the circle distributions in Figs 5A and B. First we see that the mean cellular deformation does not have a clear separation between arrangements of adhesions at the cell edge or within a constrained ring (Fig 5A). Although, in this particular example we see an increase in mean cellular deformation in 70% of the simulations when spots were moved inwards within a constrained ring relative to their corresponding (with the same angular distribution of spots) spot distribution at the cell periphery. There is a relatively even distribution of spots around the cell in the majority of these cases (with Var(*θ_g_*) ranging between 0.02 and 0.056 for 90% of the ring simulations resulting in a higher mean cellular deformation than their corresponding peripheral simulation). Figs 5E and F illustrate the corresponding cellular deformations. The cell with a distribution of spots around its periphery in Fig 5E has a lower mean deformation than that of the cell depicted in Fig 5F with the same angular distribution of spots but some more inwardly located. Conversely, in Figs 5C and D we see an example where the mean cellular deformation is greater in the case where spots are located at the periphery of the cell (Fig 5C). We explain this higher mean cellular deformation by the regions of particularly high deformation localised to the large gaps (reflected in the high value of Var(*θ_g_*), recall Figs 5C and D depict the distributions with highest Var(*θ_g_*) across the simulations presented here) between regions of adhesion in this distribution of spots.

Interestingly, we see a clear separation in the effects of the two arrangements on *W_S_* when some adhesions are distributed within a ring at the cell edge, Fig 5B. However, this is not uniformly the case, for example, when considering the same distributions of spots on a stiffer substrate (*γ* = 15) we find that 20% of the ring simulations result in a higher *W_S_* than their corresponding circle simulation, however, for the significant majority of simulations *W_S_* is reduced for the ring arrangement, see S5 Fig.

## Discussion

It is now generally accepted that the stiffness of the surrounding cell environment plays a crucial role in controlling and coordinating a range of cellular behaviours. In this study, we have shown in fact that the way in which a cell senses its environment depends critically on how the cell is adhered and not just on the mechanical stiffness of the gel. We use a model based on continuum elasticity with an active stress component to capture cellular contractility, a popular modelling framework for describing contractile cells adhered to substrates [40, 42, 46]. We here extended this model to describe spatial patterns of adhesion, considering two particular classes of cell adhesion patterns. In the first the cell is assumed to be adhered in a annular region at the cell edge, while we model the formation of small patches of adhesion, the experimentally observed focal adhesions (FAs), by the introduction of spots of adhesion distributed under the cell primarily near the cell edge. In the case of the adhered ring, analytical solutions are possible and presented, whereas the symmetry breaking inherent in the introduction of adhesive spots necessitates solutions by finite elements.

With a uniform ring of adhesion we show that the mean deformation, a measure commonly experimentally measured [48], increases with a decrease in adhered area (Fig 1). In this way, an entirely different gel stiffness can generate the same mean deformation if the adhesion percentage is carefully tuned. Similar equivalences across different stiffness have been demonstrated across other measures including maximal cellular deformation, mean cellular strain and maximal cellular strain, see S2 Fig. This observation leads us to define an effective stiffness for the combined system of the cell adhered to the gel, which quantifies the stiffness experienced by the cell. We demonstrated that the effective stiffness is always less than the true gel stiffness Table 1. The difference between the true stiffness and the effective stiffness was also shown to be reduced on stiffer substrates or arrangements with greater adhered area. When considering adhesive spots (FAs) we demonstrated that both increasing the number of spots and total adhered area through increasing spot size effectively ‘stiffens’ the substrate (Fig 2). Where the points of adhesion are randomly distributed the cell shapes qualitatively look more realistic (Fig 5C-F) and the introduction of variation in the inter-spot spacing further ‘softens’ the surface (Fig 5A).

We considered further the substrate strain energy, i.e. the work the cell does to the underlying substrate, for both adhesive ring and spots. We showed that where adhesive spots are distributed in a region around the cell edge this is energetically favourable compared with maintaining adhesions directly at the cell edge (Fig 5B), while the mean deformation is also increased, at least for the more regular arrangements (Fig 5A). We note that it is observed experimentally that FAs form on stiff substrates with soft substrates having no stable adhesions, and that FAs grow and elongate on these substrates [15,20]. Significantly, we here show that starting with nascent adhesions of small area, FA growth and elongation would be energetically favourable on stiff substrates but not on soft substrates (Figs 4D and E). We additionally show that although FA growth would by itself reduce the mean deformation and effectively ‘stiffen’ the substrate the elongation of the adhesion spot can compensate for this effect (Fig 3). On soft substrates spot elongation also reduces *W_s_*. However, on stiff substrates spot elongation only reduces the work done up to an optimal elongation. Indeed we predict an optimal spot elongation of 2-3 for e.g. *γ* = 15 across a range of adhesion profiles. A resistance parameter of *γ* =15 corresponds to a Young’s modulus for the substrate of *E_S_* ≈ 318kPa, with cell parameters *E_c_* = 10kPa, radius *r*_0_ = 30*μ*m, thickness *h_c_* = 1*μ*m, cell and substrate Poisson’s ratios *ν_c_* = *ν_S_* = 0.45 and substrate thickness *h_S_* = 35*μ*m [48,53]. We thus suggest an underpinning mechanism driving the observed FA dynamics based on energy considerations.

Taken together our results indicate that mechanotransduction studies require a consideration of the combined cell and substrate system moving beyond a focus on individual FAs. It is clear that there is a need for integrating theoretical modelling with experimental investigations to enable the full complexity of the system to be accounted for. However, taking this forward into mechanotransduction studies is non-trivial. Although fluorescence imaging and segmentation of FAs is a well-established technique, it is still technically complex and the adhesion patterns vary greatly even between cells on the same surface. In this context, studies in which cell adhesion is directly controlled through micropatterning techniques [21, 22] could have significant advantages; where the adhesion patterns are determined *a priori* this can be controlled for across experiments.

## Methods

### Cell mechanical model and analytical solution for adhesive ring

The mechanical model laid out above is based on an active stress model for cell contractility. Thus the cell stress is considered to comprise of two components such that *σ* = *σ^P^* + *σ^A^*, where 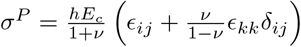 is the constitutive relation between stress and strain for a linearly elastic material in plane stress and 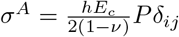, is the active component representing active contraction analogous to thermal cooling where *P* is the target area reduction of cell elements [40]. We take *P* = −*P*_0_ constant throughout. In these constitutive relations, *E_c_* and *ν* are the cell’s Young’s modulus and Poisson’s ratio respectively, *δ_ij_* the Kronecker delta and *ϵ_ij_* the linear strain tensor. Linear elasticity is a common simplifying assumption for small strains [20,26,40–42] with the timescales over which cell adhesion, contractility and sensing occurring being faster than those of viscoelastic relaxation justifying an elastic assumption [42]. Throughout, we impose the condition that there is zero stress at the outer cell edge.

For the case where the cell is assumed to adhere in a ring symmetry ensures that all deformations are radial, i.e. the deformation **u** = *u*(*r*)**e**_*r*_. On substituting the constitutive relations into the force balance (1) we obtain

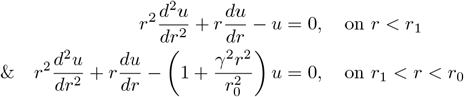

where the cell is adhered in *r*_1_ < *r* < *r*_0_, which has exact solution (2) with the constants *α*_0_, *α*_1_, *β*_1_ given by

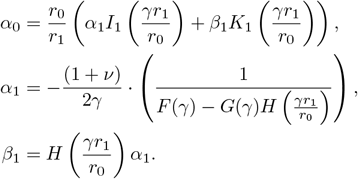

For conciseness we introduced functions:

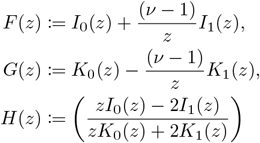

In the limit 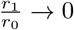, we recover the solution of a completely adhered disc [40]

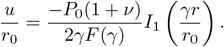

The mean cellular deformation of a cell with an adhered ring can be calculated as

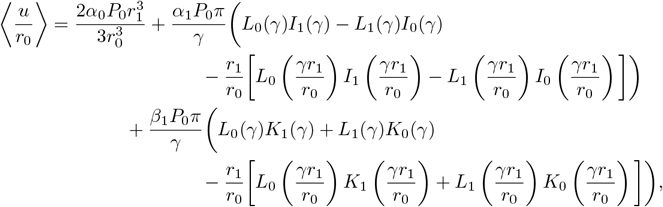

where *L*_0_ and *L*_1_ are modified Struve functions [54].

From the solution for u the strain energy *W* = *W_CA_* + *W_CP_* + *W_S_* can be calculated from the integral expressions

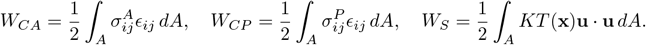

With the approximation of the substrate as a linear array of springs there are no non-local effects with no deformation outside of the cell spread area *A* so the integrals can be restricted to *A* without loss of generality. We note that minimising the energy functional *W* is equivalent to solving the force balance equation (1) [26,42,55] (although [26, 55] include an additional term to account for the deformation of the cell membrane). We scale all energies by a factor of (1 – *ν*^2^)/*hE_c_r*_0_^2^.

The energies are calculated numerically for the case of the adhered spots, however, for a adhered disc analytical solutions are possible. For example, for the adhesive ring the substrate strain energy *W_S_* can be analytically calculated (using the known integreal expressions for Bessel functions [54]) as

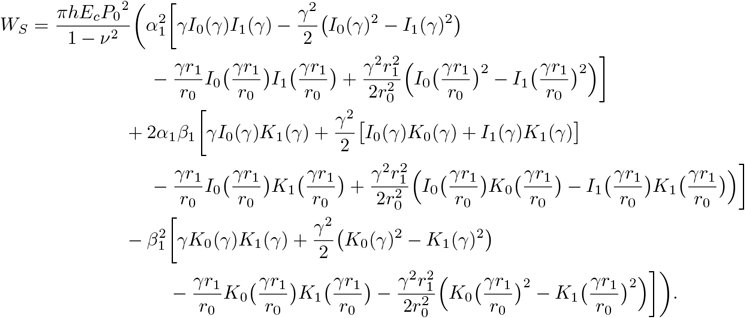

### Numerical solutions for adhered spots

Where the cell is assumed to be adhered to the substrate in spots the symmetry of the problem is broken and numerical methods must be used. All solutions were obtained using finite element methods in MATLAB (PDE Toolbox R2018a). The cell geometry was specified as in Fig 2A, for example, and a triangular mesh generated. In order to calculate the integral for the mean cellular deformation and energy integrals from the numerically computed data the Gaussian quadrature of degree 1 was used to approximate the solution on each triangle of the mesh. For a regular arrangement of spots circles of adhesion were placed at angles 2*π*/*N*, where *N* is the number of spots; the spots are placed radially at a position 0.98*r*_0_ – *r_s_*, where *r_s_* is the spot radius. In order to simulate non-uniform spot distributions we consider two cases. In both instances we considered 20 spots, each the same size and totalling an adhered area of 10% of the cell spread area. First, we distributed the spots around the edge of a cell (assumed to be circular with radius *r*_0_), dividing a circle of radius 0.91*r*_0_ into 37 ‘bins’ such that each was large enough for 1 spot with no overlaps. In each of our simulations, 20 bins were randomly sampled, resulting in a distribution of 20 spots around the cell edge (eg. Fig 5C and E). Second, we considered spots distributed in an annular region located near the cell edge. In order for direct comparison of the effect of variation in the radial position of the spots as opposed to located at the edge, we considered the same angular distributions of spots as in the edge simulations. The radial position of the centre of each spot was randomly sampled from a uniform distribution between 0.6*r*_0_ and 0.91*r*_0_. The distance between adjacent spots was tested to ensure there was no overlap. In cases where the simulations did result in distributions with overlapping spots, the radial position of spot to the ‘left’ was re-sampled until there was no overlap.

## Supporting information

**S1 Fig. On stiffer substrates, cells require a thinner ring to achieve the same proportion of effectively experienced resistance.** Heat map shows relative effective resistance plotted against internal ring radius (*r*_1_/*r*_0_) and *γ*; *ν* = 0.45 and *P*_0_ = 0.7.

**S2 Fig. Other quantitative measures demonstrate similar effects to mean cellular deformation.** Heat maps show (A) the maximum cellular deformation, (B) maximum cellular strain, and (C) mean cellular strain, plotted against ring thickness (parameterised by *r*_1_) and substrate stiffness (parameterised by *γ*). For each postulated measure, we would predict the same effective resistance experienced along the contour lines (black) which show a constant value of the given measure. These appear qualitatively very similar across the plots.

**S3 Fig. W_CA_ dynamics are qualitatively very similar to the mean cellular deformation.** Active work done by the cell plotted against adhered area for cells with different numbers of evenly distributed adhesive spots; *γ* = 7, *ν* = 0.45, *P*_0_ = 0.7.

**S4 Fig. Focal Adhesion growth reduces strain energy on stiff substrates.** Heat map shows substrate strain energy resulting from a cell with an adhered ring against inner ring radius (*r*_1_/*r*_0_) and substrate stiffness (*γ*).

**S5 Fig. Random placement of adhesion sites for stiff substrates with *γ* = 15.** (A) Mean cellular deformation, and (B) substrate strain energy, plotted against the variance in angular gap size between adjacent spots from 20 simulations of randomly placed spots around a cell edge and corresponding ‘ring’ distributions with the same angular spot placements but radial positions distributed within an annular region between 0.6r0 and the cell edge. Spot distributions are identical to those in Figs 5A and B. Results for an even distribution of spots are included for comparison.

## Acknowledgments

JSW acknowledges PhD funding from the UK Engineering and Physical Sciences Research Council, institutional Doctoral Training Partnership (grant EP/N509772/1). CMD also acknowledges financial support from the UK EPSRC (grant EP/M012964/1).

## References

1. Vogel V, Sheetz M. Local force and geometry sensing regulate cell functions. Nat Rev Mol Cell Biol. 2006;7(4):265–275.

2. Wozniak MA, Chen CS. Mechanotransduction in development: A growing role for contractility. Nat Rev Mol Cell Biol. 2009;10(1):34–43.

3. Heer NC, Martin AC. Tension, contraction and tissue morphogenesis. Development. 2017;144(23):4249–4260.

4. Irvine KD, Shraiman BI. Mechanical control of growth: Ideas, facts and challenges. Development. 2017;144(23):4238–4248.

5. Wolfenson H, Yang B, Sheetz MP. Steps in mechanotransduction pathways that control cell morphology. Annu Rev Physiol. 2019;81:585–605.

6. Engler AJ, Sen S, Sweeney HL, Discher DE. Matrix elasticity directs stem cell lineage specification. Cell. 2006;126(4):677–689.

7. Kumar A, Placone JK, Engler AJ. Understanding the extracellular forces that determine cell fate and maintenance. Development. 2017;144(23):4261–4270.

8. Verbruggen SW, Vaughan TJ, McNamara LM. Mechanisms of osteocyte stimulation in osteoporosis. J Mech Behav Biomed. 2016;62:158–168.

9. Segel M, Neumann B, Hill MF, Weber IP, Viscomi C, Zhao C, et al. Niche stiffness underlies the ageing of central nervous system progenitor cells. Nature. 2019;573(7772):130–134.

10. Polacheck WJ, Chen CS. Measuring cell-generated forces: a guide to the available tools. Nat Methods. 2016;13(5):415–423.

11. Roca-Cusachs P, Conte V, Trepat X. Quantifying forces in cell biology. Nat Cell Biol. 2017;19(7):742–751.

12. Missirlis D, Spatz JP. Combined effects of PEG hydrogel elasticity and cell-adhesive coating on fibroblast adhesion and persistent migration. Biomacromolecules. 2014;15(1):195–205.

13. Schwarz US, Bischofs IB. Physical determinants of cell organization in soft media. Med Eng Phys. 2005;27(9):763–772.

14. Discher DE, Janmey P, Wang Y. Tissue cells feel and respond to the stiffness of their substrate. Science. 2005;310(5751):1139–1143.

15. Geiger B, Spatz JP, Bershadsky AD. Environmental sensing through focal adhesions. Nat Rev Mol Cell Biol. 2009;10(1):21–23.

16. Mullen CA, Vaughan TJ, Voisin MC, Brennan MA, Layrolle P, McNamara LM. Cell morphology and focal adhesion location alters internal cell stress. J R Soc Interface. 2014;11(101):20140885.

17. McKenzie AJ, Hicks SR, Svec KV, Naughton H, Edmunds ZL, Howe AK. The mechanical microenvironment regulates ovarian cancer cell morphology, migration, and spheroid disaggregation. Sci Rep. 2018;8(1):1–20.

18. Ghibaudo M, Saez A, Trichet L, Xayaphoummine A, Browaeys J, Silberzan P, et al. Traction forces and rigidity sensing regulate cell functions. Soft Matter. 2008;4(9):1836–1843.

19. Oakes PW, Gardel ML. Stressing the limits of focal adhesion mechanosensitivity. Curr Opin Cell Biol. 2014;30:68–73.

20. Schwarz US, Safran SA. Physics of adherent cells. Rev Mod Phys. 2013;85(3):1327.

21. Liu WF, Chen CS. Cellular and multicellular form and function. Adv Drug Deliv Rev. 2007;59(13):1319–1328.

22. Théry M. Micropatterning as a tool to decipher cell morphogenesis and functions. J Cell Sci. 2010;123(24):4201–4213.

23. McWhorter FY, Wang T, Nguyen P, Chung T, Liu WF. Modulation of macrophage phenotype by cell shape. Proc Natl Acad Sci. 2013;110(43):17253–17258.

24. Jain N, Iyer KV, Kumar A, Shivashankar GV. Cell geometric constraints induce modular gene-expression patterns via redistribution of HDAC3 regulated by actomyosin contractility. Proc Natl Acad Sci. 2013;110(28):11349–11354.

25. Chen CS, Mrksich M, Huang S, Whitesides GM, Ingber DE. Geometric control of cell life and death. Science. 1997;276(5317):1425–1428.

26. Oakes PW, Banerjee S, Marchetti MC, Gardel ML. Geometry regulates traction stresses in adherent cells. Biophys J. 2014;107(4):825–833.

27. Charrier EE, Pogoda K, Wells RG, Janmey PA. Control of cell morphology and differentiation by substrates with independently tunable elasticity and viscous dissipation. Nat comms. 2018;9(1):1–13.

28. Iskratsch T, Wolfenson H, Sheetz MP. Appreciating force and shape—the rise of mechanotransduction in cell biology. Nat Rev Mol Cell Biol. 2014;15(12):825–833.

29. Fletcher DA, Mullins RD. Cell mechanics and the cytoskeleton. Nature. 2010;463(7280):485–492.

30. Schwarz US, Gardel ML. United we stand–integrating the actin cytoskeleton and cell-matrix adhesions in cellular mechanotransduction. J Cell Sci. 2012;125(13):3051–3060.

31. Gardel ML, Kasza KE, Brangwynne CP, Liu J, Weitz DA. Mechanical response of cytoskeletal networks. Methods Cell Biol. 2008;89:487–519.

32. Riveline D, Zamir E, Balaban NQ, Schwarz US, Ishizaki T, Narumiya S, et al. Focal contacts as mechanosensors: externally applied local mechanical force induces growth of focal contacts by an mDia1-dependent and ROCK-independent mechanism. J Cell Biol. 2001;153(6):1175–1186.

33. Galbraith CG, Yamada KM, Sheetz MP. The relationship between force and focal complex development. J Cell Biol. 2002;159(4):695–705.

34. Balaban NQ, Schwarz US, Riveline D, Goichberg P, Tzur G, Sabanay I, et al. Force and focal adhesion assembly: a close relationship studied using elastic micropatterned substrates. Nat Cell Biol. 2001;3(5):466–472.

35. Cho S, Irianto J, Discher DE. Mechanosensing by the nucleus: From pathways to scaling relationships. J Cell Biol. 2017;216(2):305–315.

36. Song Y, Soto J, Chen B, Yang L, Li S. Cell engineering: Biophysical regulation of the nucleus. Biomaterials. 2020;234:119743.

37. Albert PJ, Schwarz US. Modeling cell shape and dynamics on micropatterns. Cell Adh Migr. 2016;10(5):516–528.

38. Freedman SL, Banerjee S, Hocky GM, Dinner AR. A versatile framework for simulating the dynamic mechanical structure of cytoskeletal networks. Biophys J. 2017;113(2):448–460.

39. Shishvan SS, Vigliotti A, Deshpande VS. The homeostatic ensemble for cells. Biomech Model Mechanobiol. 2018;17(6):1631–1662.

40. Edwards CM, Schwarz US. Force localization in contracting cell layers. Phys Rev Lett. 2011;107(12):128101.

41. Banerjee S, Marchetti MC. Substrate rigidity deforms and polarizes active gels. Europhys Lett. 2011;96(2):28003.

42. Friedrich BM, Safran SA. How cells feel their substrate: spontaneous symmetry breaking of active surface stresses. Soft Matter. 2012;8(11):3223–3230.

43. Kohn JC, Abdalrahman T, Sack KL, Reinhart-King CA, Franz T. Cell focal adhesion clustering leads to decreased and homogenized basal strains. Int J Numer Meth Bio. 2019;35(12):e3260.

44. Dunlop C. Differential cellular contractility as a mechanism for stiffness sensing. N J Phys. 2019;21(6):063005.

45. Banerjee S, Marchetti MC. Contractile stresses in cohesive cell layers on finite-thickness substrates. Phys Rev Lett. 2012;109(10):108101.

46. Banerjee S, Marchetti MC. Controlling cell-matrix traction forces by extracellular geometry. New J Phys. 2013;15(3):035015.

47. Tan JL, Tien J, Pirone DM, Gray D, Bhadriraju K, Chen CS. Cells lying on a bed of microneedles: an approach to isolate mechanical force. Proc Natl Acad Sci USA. 2003;100(4):1484–1489.

48. Saez A, Anon E, Ghibaudo M, du Roure O, Di Meglio JM, Hersen P, et al. Traction forces exerted by epithelial cell sheets. J Phys Condens Matter. 2010;22(19):194119.

49. Butler JP, Tolic-Nørrelykke IM, Fabry B, Fredberg JJ. Traction fields, moments, and strain energy that cells exert on their surroundings. Am J Physiol Cell Physiol. 2002;282(3):C595–C605.

50. Koch TM, Munster S, Bonakdar N, Butler JP, Fabry B. 3D traction forces in cancer cell invasion. PloS one. 2012;7(3):e33476.

51. Mierke CT, Frey B, Fellner M, Herrmann M, Fabry B. Integrin α5β1 facilitates cancer cell invasion through enhanced contractile forces. J Cell Sci. 2011;124(3):369–383.

52. Oakes PW. Balancing forces in migration. Curr Opin Cell Biol. 2018;54:43–49.

53. Prager-Khoutorsky M, Lichtenstein A, Krishnan R, Rajendran K, Mayo A, Kam Z, et al. Fibroblast polarization is a matrix-rigidity-dependent process controlled by focal adhesion mechanosensing. Nat Cell Biol. 2011;13(12):1457–1465.

54. Abramowitz M, Stegun IA. Handbook of mathematical functions: with formulas, graphs, and mathematical tables. vol. 55. Courier Corporation; 1964.

55. Banerjee S, Sknepnek R, Marchetti MC. Optimal shapes and stresses of adherent cells on patterned substrates. Soft Matter. 2014;10(14):2424–2430.

